# Altered brain structure in an ATRX-deficient mouse model of autism spectrum disorder

**DOI:** 10.1101/2025.08.06.666605

**Authors:** Katherine Quesnel, Jacob Ellegood, Jason P. Lerch, Nathalie G. Bérubé

**Author notes:** **Authors contact:** Katherine Quesnel contact, Jacob Ellegood contact Jason P Lerch contact.

## Abstract

Mutations in the *ATRX* gene are a primary cause of ATR-X syndrome, which is characterized by intellectual disability, autism, and a range of brain structural abnormalities, including microcephaly. We previously showed that mice with conditional ATRX ablation in forebrain excitatory neurons display deficits in fear memory and autism-related behaviors, with some effects exhibiting sexual dimorphism. In this study, we used high-resolution magnetic resonance imaging (MRI) to systematically characterize brain structural changes associated with these behavioral abnormalities. Whole-brain analysis revealed male-specific microcephaly, while subregional analysis identified significant reductions in hippocampal structures and increased volume of the caudal cortex in mutant animals of both sexes. We also identified structural alterations in regions retaining ATRX expression, such as the thalamus, midbrain, cerebellum, and several fiber tracts. These findings suggest that ATRX loss disrupts the coordinated development of interconnected brain regions. Overall, our results implicate impaired cortico-thalamic-cerebellar connectivity as a potential neural substrate underlying the autistic-like behaviors observed in this mouse model, providing new insights into the neurobiological basis of ATR-X syndrome.

**LAY SUMMARY:** Changes in a gene called ATRX are known to affect brain development and are linked to intellectual disability and autism. In our previous work, we found that removing this gene early in brain development caused mice to show behaviors like those seen in people with autism. In this study, we used detailed brain scans to see if these behavioral changes were linked to differences in brain structure. We found that male mice without ATRX had smaller brains and bodies, while female mice did not show the same brain size reduction. However, both male and female mice had smaller areas of the brain important for memory and movement, and larger areas involved in thinking and sensing. We also saw changes in parts of the brain where ATRX was still present, suggesting that early changes in one area can affect how the whole brain develops. These findings help us understand how early disruptions in brain development might lead to autism-related behaviors.

## INTRODUCTION

Alpha-thalassemia intellectual disability X-linked (ATRX), is a large chromatin remodeling protein crucial for genomic integrity and maintenance of heterochromatin (Bérubé et al., 2000; Gibbons & Higgs, 2000). Hypomorphic mutations in *ATRX* are associated with ATR-X syndrome which is characterized by mild-to-severe intellectual disability (ID), as well as non-syndromic ID. *ATRX* is also classified as high-risk gene for autism spectrum disorder (ASD). Patients with ATR-X syndrome exhibit global neurodevelopment delays, skeletal malformations such as short stature and craniofacial alterations, and a portion of patients are reported to have microcephaly (Gibbons et al., 2008; Gibbons & Higgs, 2000).

We have previously characterized a mouse model with early embryonic deletion of *Atrx* in excitatory neurons of the forebrain. In this model, both male and female adult mice display fear memory impairments, hyperactivity and repetitive behaviours– a core ASD feature. Additionally, male mice specifically display auditory sensory gating deficits, social memory deficits and social aggression (Quesnel et al., 2023). Further characterization of this model will help elucidate the developmental consequences of ATRX loss, including the specific brain structures affected and how these alterations contribute to the ASD features observed.

ASD is highly heterogeneous, both in terms of the phenotypes displayed and the causative genes, many of which are unknown. Some studies have tried to cluster ASD patients according to MRI findings based on neuroanatomical abnormalities relating to their clinical features (Hong et al., 2018; Liu et al., 2022). Investigating mouse models with specific behavioural phenotypes can help directly link brain structural alterations to corresponding behaviours. For instance, a study examining 26 different ASD mouse models found common brain volume changes in regions such as the cerebellar cortex and striatum—areas frequently implicated in human ASD MRI studies (Ellegood et al., 2015). A recent study identified consistent neuroanatomical clusters in both mouse models and human NDD populations, linking brain structure to underlying molecular pathways and highlighting connections between gene interactions, neuroanatomy, and behavioral phenotypes (Ellegood et al. 2025). The cerebellum has been shown to be disrupted in mouse models displaying social deficits (Al Sagheer et al., 2018; Haida et al., 2019), while striatal volume in ASD patients has been correlated with repetitive behaviors (Hollander et al., 2005). In our study, we aim to determine whether similar neuroanatomical changes arise from the loss of ATRX, and whether these changes are related to the ASD-like behaviors previously observed. This will help elucidate the impact of ATRX loss during brain development on adult brain structure.

ATR-X syndrome is rare, with just over 200 reported cases, and exhibits a range of neurodevelopmental phenotypes. An MRI study of 27 ATR-X patients revealed considerable variability in neuroanatomical alterations (Wada et al., 2013). Notably, one patient showed underdevelopment of the cerebellar vermis, and four patients had underdevelopment of the corpus callosum, supporting previous findings that the cerebellum and disrupted connectivity play a role in ASD (Ellegood et al., 2015). Another mouse model with post-natal deletion of *Atrx* in forebrain excitatory neurons displayed alterations in hippocampal regions correlating with memory deficits, but no ASD-features were detected in this model (Martin-Kenny & Bérubé, 2020; Tamming et al., 2020). Therefore, comparing our earlier timepoint for *Atrx* deletion with the presence of ASD-features could identify brain regions correlating with memory deficits vs ASD associated behaviours. Our findings indicate significant reduction in brain size, decreased relative volumes in the hippocampus and cerebellum and their associated fiber tracts, alongside enlargement of caudal cortical subregions, thalamus, and midbrain regions. These results suggest that disruption in cortico-thalamic-cerebellar pathways may contribute to the ASD behaviours characteristic of this mouse model.

## MATERIALS AND METHODS

### Animal care and husbandry

Conditional inactivation of *Atrx* in post-mitotic excitatory neurons was achieved beginning at embryonic day 11.5 (E11.5). To generate experimental animals, 129SV female mice heterozygous for *Atrx*^loxP^ alleles (Berube et al., 2005), were crossed with C57BL/6J male mice expressing Cre recombinase under the control of the NEX-Cre promoter (Goebbels et al., 2006). Male offspring carrying a floxed *Atrx* allele on the X chromosome and the NEX-Cre transgene (*Atrx*^NEXCre^ males) served as knockout animals, while male littermates with a wild-type *Atrx* allele and the NEX-Cre transgene were used as controls to account for any effects of Cre expression. Similarly, homozygous *Atrx* floxed female mice expressing NEX-Cre (*Atrx*^NEXCre^ females) were used as female knockouts, with wild-type *Atrx*, NEX-Cre-positive females serving as controls. Genotyping was performed as previously described (Quesnel et al., 2023).

To generate Cre-reporter mice, Sun1-GFP mice [B6;129-Gt(ROSA)26Sortm5(CAG-Sun1/sfGFP)Nat/J; IMSR JAX:021039; MGI:5614796] (Mo et al., 2015) were crossed with C57BL/6 *Atrx*^loxP^ mice. This cross produced C57BL/6 female offspring heterozygous for both the *Atrx*^loxP^ allele and the Sun1-GFP reporter. These double heterozygous females were then incorporated into the breeding scheme described above to enable tracing of brain regions with Cre-mediated recombination and subsequent *Atrx* ablation. All animal procedures were conducted in accordance with the regulations of the Animals for Research Act of the province of Ontario and approved by the University of Western Ontario Animal Care and Use Committee (protocols 2021-049 and 2021-064). Mice were housed under a 12-hour light/12-hour dark cycle with ad libitum access to food and water.

### Immunofluorescence staining

Mice 3-months of age (n=4) were perfused transcardially with cold 1× PBS, followed by cold 4% paraformaldehyde (PFA). Brains were post-fixed overnight in 4% PFA at 4°C, then washed in 1× PBS and sunk in 30% sucrose for a minimum of one week. Tissues were embedded in Cryomatrix™ (Epredia, cat# 6769006) using an ethanol–dry ice slurry and stored at −80°C. Brains were sectioned at 10 μm thickness using a Leica CM3050 S cryostat and mounted onto Superfrost Plus microscope slides (Fisher Scientific, cat# 12-550-15). Sections were rehydrated in 1× PBS and subjected to antigen retrieval by boiling in sodium citrate buffer (pH 6.0, 0.5% Tween-20) for 15 minutes. Permeabilization was performed with 1× PBS containing 0.03% Triton X-100 for 10 minutes, followed by blocking for 1 hour in 1× PBS with 0.3% Triton X-100 and 5% goat serum. Primary antibodies diluted in blocking solution were applied overnight at 4°C: anti-Atrx (Santa Cruz, sc15408; rabbit, 1:150) and anti-GFP (Invitrogen, PA1-95333; chicken, 1:200). After washing with 1× PBS containing 0.3% Triton X-100, sections were incubated for 1 hour at room temperature with the appropriate secondary antibodies (Alexa Fluor 594 anti-rabbit, Thermo Scientific, cat# A11012, 1:1000; Alexa Fluor 488 anti-chicken, Thermo Scientific, cat# A11039, 1:1000). Slides were washed, counterstained with DAPI, rinsed, and coverslipped using ImmunoMount (Thermo Scientific, cat# 9990402). Images were acquired using a Leica CTR 6500 microscope with OpenLab and Velocity software.

### Brain histological analysis

Coronal brain sections were stained with hematoxylin and eosin (H&E) to assess overall brain structure. Sections were first rehydrated in 70% ethanol for 5 minutes, followed by a 5-minute rinse in distilled water. Slides were then immersed in hematoxylin solution for 2 minutes, rinsed under cold running tap water for 30 seconds, incubated in Tasha’s bluing solution for 30 seconds, and washed in tap water for 8 minutes. Sections were subsequently stained with 5% eosin Y for 2 minutes, followed immediately by two washes in 70% ethanol. Dehydration was performed with a 1-minute wash in 90% ethanol, followed by two 2-minute washes in 100% ethanol. Sections were then cleared in xylene with three 5-minute washes and mounted using Permount Mounting Media (Fisher Chemical, SP15-100). Two weeks post-staining, slides were cleaned and imaged using a Leica Aperio ScanScope. Images were analyzed with ImageScope software. Quantification of total brain area (mm²) and corpus callosum thickness (mm) was performed using ImageJ software.

### Magnetic resonance ex-vivo brain imaging

Mice 10-13 months of age (Ctrl^male^ n=10, *Atrx*^NEXCre^ ^male^ n=11, Ctrl^female^ n=16, *Atrx*^NEXCre^ ^female^ n=12) were anesthetized with a mixture of ketamine/xylazine and intracardially perfused with 30mL of 1xPBS containing 0.05 U/mL heparin (Sigma) and 2mM ProHance (Bracco Diagnostics, a Gadolinium contrast agent) followed by 30 mL of 4% PFA containing 2mM ProHance (Cahill et al., 2012; Lerch et al., 2011). Perfusions were performed at a rate of approximately 2 mL/min. After perfusion, mice were decapitated and the skin, lower jaw, ears, and the cartilaginous nose tip were removed. The brain and remaining skull structures were incubated in 4% PFA + 2mM ProHance overnight at 4°C then transferred to 1xPBS containing 2mM ProHance and 0.02% sodium azide for at least 1 month prior to scanning (de Guzman et al., 2016).

A 7-Tesla 306 mm horizontal bore magnet (BioSpec 70/30 USR, Bruker, Ettlingen, Germany) with a ParaVision 6.0.1 console was used to image brains within their skull. Eight samples were imaged in parallel using a custom-built 8-coil solenoid array. To acquire anatomical images, the following scan parameters were used: T2-weighted 3D fast spin echo (FSE) sequence with a cylindrical acquisition of k-space, TR/TE/ETL = 350 ms/12 ms/6, TEeff = 30ms, four effective averages, FOV/matrix-size = 20.2 × 20.2 × 25.2 mm / 504 × 504 × 630, total-imaging-time = 13.2 h. The resulting anatomical images had an isotropic resolution of 40µm voxels.

### Image registration and analysis

Registration consisted of both linear (rigid then affine) transformations and non-linear transformations. These registrations were performed with a combination of mni_autoreg tools (Collins et al., 1994) and ANTS (advanced normalization tools) (Avants et al., 2008, 2011). After registration, all scans were resampled with the appropriate transform and averaged to create a population atlas representing the average anatomy of the study sample. The result of these registrations were deformation fields that transform images to a consensus average. Therefore, these deformations fields quantify anatomical differences between images. As detailed in previous studies (Lerch et al., 2008; Nieman et al., 2006), the Jacobian determinants of the deformation fields were computed and analyzed to measure the volume differences between subjects at every voxel. A pre-existing classified MRI atlas was warped onto the population atlas (containing 282 different segmented structures encompassing cortical lobes, large white matter structures such as the corpus callosum, ventricles, cerebellum, brain stem, and olfactory bulbs (Dorr et al., 2008; Qiu et al., 2018; Richards et al., 2011; Steadman et al., 2014; Ullmann et al., 2013) to compute the volume of brain structures in all the input images. A linear model with a genotype predictor was used to assess significance. The model was either fit to the volume of every structure independently (structure-wise statistics) of fit to every voxel independently (voxel-wise statistics), and multiple comparisons in this study were controlled for using the False Discovery Rate (Genovese et al., 2002). FDR <0.10 was considered statistically significant.

## RESULTS

### Sex-specific microcephaly and regional brain alterations in *Atrx*^NEXCre^ mice

Ex vivo MRI analysis of 12-month-old mice revealed significant microcephaly in *Atrx*^NEXCre^ compared control male mice (FDR = 0.0065), whereas female *Atrx*^NEXCre^ mice exhibited no global brain volume differences (**Figure 1A**). This male-specific reduction in brain size parallels the heightened neurological severity observed in the mutant male mice (Quesnel et al., 2023) and aligns with the male predominance of autism spectrum disorder (ASD) diagnoses. Both male and female *Atrx*^NEXCre^ mice displayed reduced body size at 12 months, including decreased body weight (males: *p* < 0.0001; females: *p* = 0.0162) and shorter nose-to-tail length (males: *p* = 0.0091; females: *p* = 0.0426) (**Figure 1B**). Brain size strongly correlated with body size in males (*r* = 0.82, *p* = 0.001) but not in female mutant mice (*r* = 0.24, *p* = 0.38), suggesting sex-specific mechanisms governing neurodevelopmental scaling.

**Figure 1:**
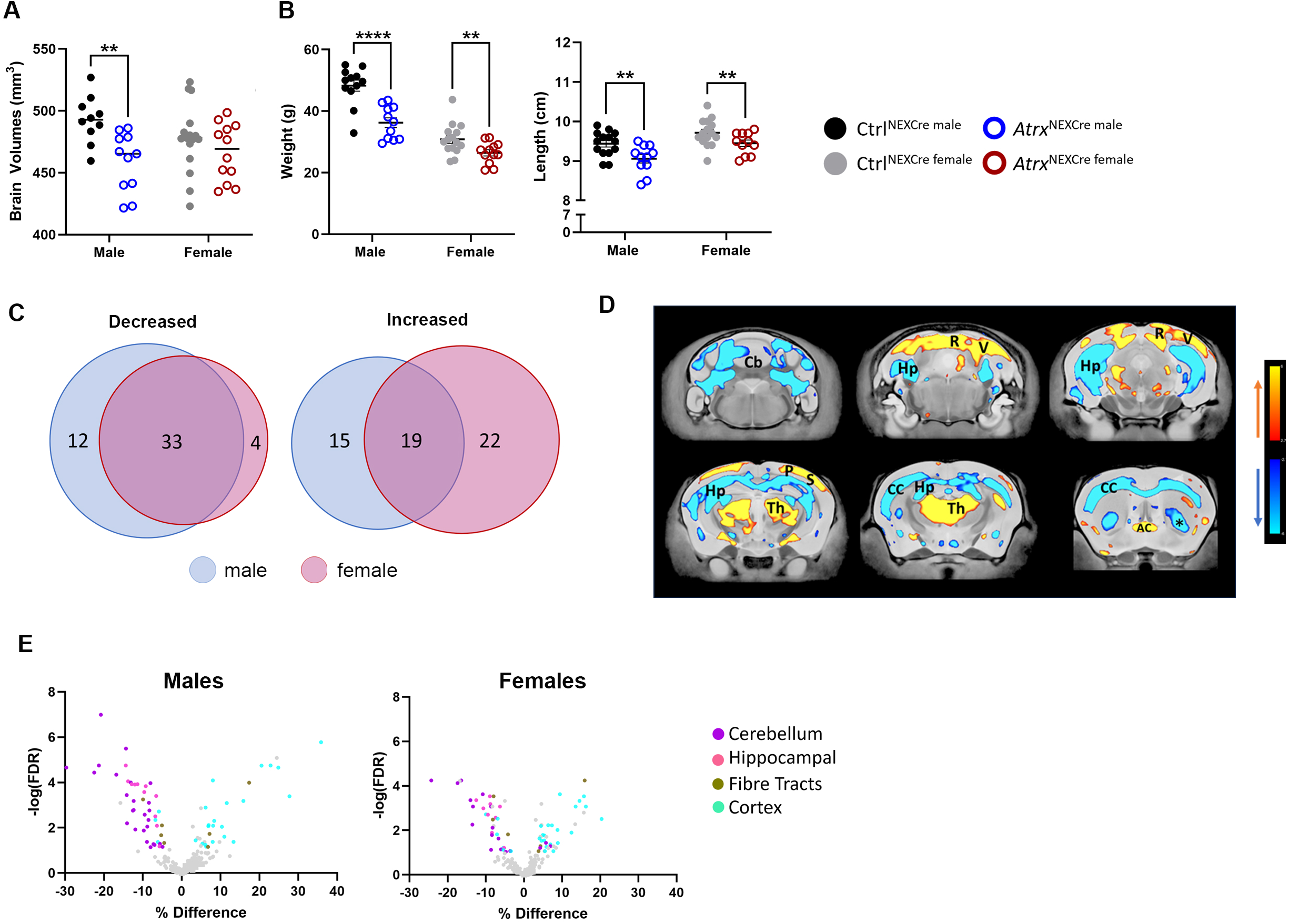
Reproducible regional brain volumetric changes in the *Atrx*^NEXCre^ compared to control mice. **A)** Total brain volume and **B)** body weight and length of 12-month-old *Atrx*^NEXCre^ and control mice. Total brain volume: **FDR < 0.05. Body size: **p <0.05, ****p < 0.0001). **C)** Venn diagrams displaying the overlap of brain subregions with significantly altered relative volume in male and female *Atrx*^NEXCre^ mice. **D)** Coronal brain images highlight areas with significantly increased (red-yellow spectrum) and decreased (blue spectrum) relative volume. **E)** Volcano plot of 188 colour-coded significantly changed brain subregions [cerebellum (purple), hippocampus (pink), fibre tracts (gold), cortex (blue)]. Cb= cerebellum, Hp= Hippocampus, CC= corpus callosum, Th= thalamus, R= retrosplenial cortex, P= parietal cortex, V= visual cortex, AC= anterior commissure, *= posterior ventral caudoputamen (Ctrl^male^ n=10, *Atrx*^NEXCre male^ n=11, Ctrl^female^ n=16, *Atrx*^NEXCre female^ n=12).

To account for total brain volume differences, subsequent analyses used relative regional volumes. In males, 45 of 188 brain subregions showed decreased relative (i.e. after normalizing for overall brain size) volume, while 34 exhibited increases (FDR < 0.1). Females displayed similar patterns, with 37 subregions decreased and 41 increased (**Figure 1C,D).** Across both sexes, volume reductions predominated in the cerebellum and hippocampus—regions critical for motor coordination and memory—while expansions occurred primarily in cortical areas. Fiber tracts showed both increased and decreased relative volume (**Figure 1E**).

### Ultrastructural differences in hippocampal and cortical regions

Significant alterations in relative volume were observed in two major forebrain regions—the hippocampus and the cortex—of *Atrx*^NEXCre^ mice. The hippocampal formation exhibited a marked reduction in relative volume in both male (FDR = 0.0028) and female (FDR = 0.0041) *Atrx*^NEXCre^ brains compared to controls (**Figure 2A**). This reduction aligns with previously reported memory deficits in this model, underscoring the functional relevance of these structural changes. Further analysis of hippocampal subregions revealed that the subiculum, CA1, CA2, and CA3 regions were significantly reduced, whereas the dentate gyrus remained unaffected (**Figure 2B**).

**Figure 2:**
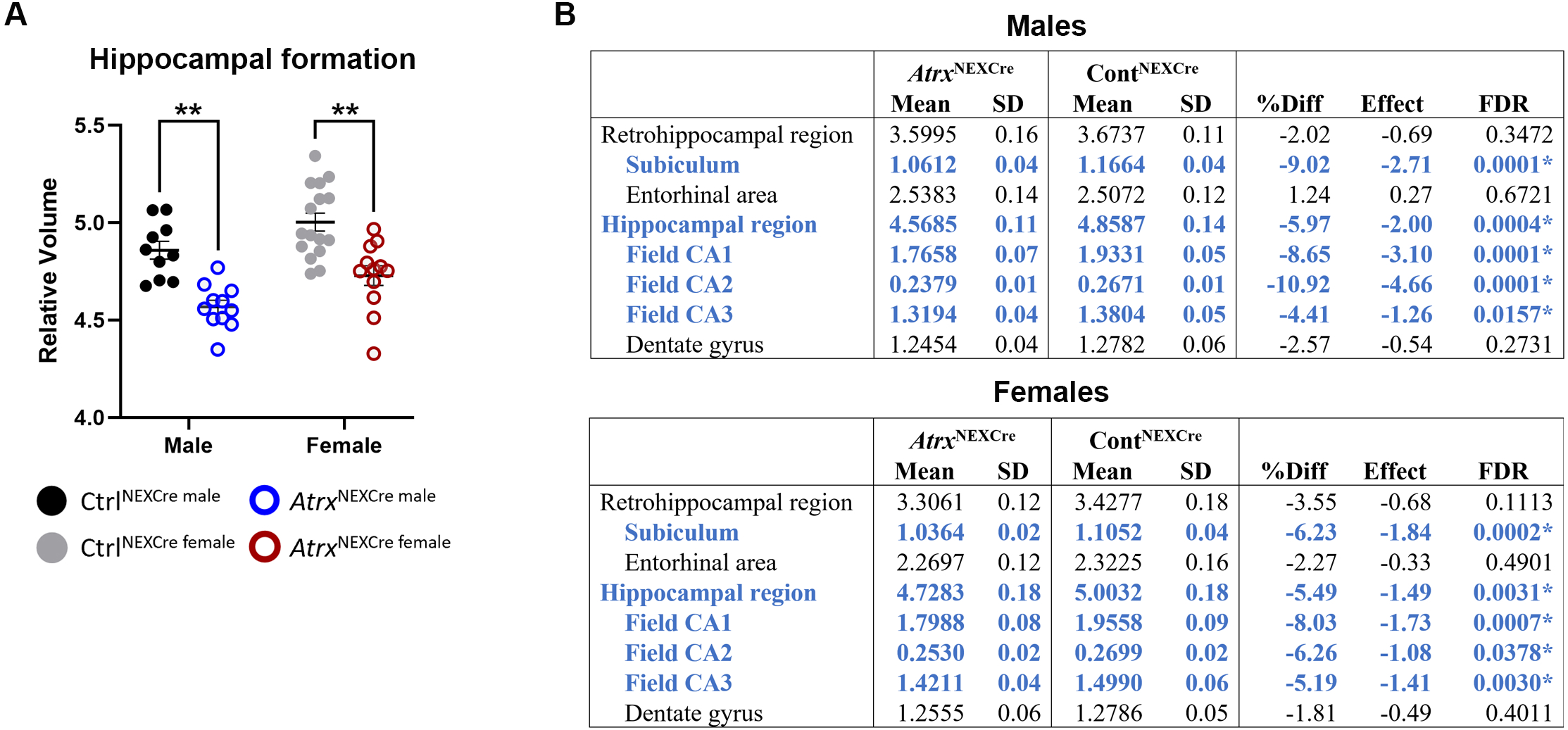
Significantly decreased relative volume of the hippocampal formation in *Atrx*^NEXCre^ mice. **A)** Decreased relative volume of the hippocampal formation and **B)** of specific hippocampal subregions for male and female mice. Blue= significant relative decrease in volume. SD: standard deviation, FDR: false discovery rate. (**= FDR <0.05) (Ctrl^male^ n=10, *Atrx*^NEXCre male^ n=11, Ctrl^female^ n=16, *Atrx*^NEXCre female^ n=12).

In contrast, the total isocortex displayed a significant increase in relative volume in male *Atrx*^NEXCre^ mice (FDR = 0.0005), an effect not observed in females (**Figure 3A**). Subregional analysis of the cortex in males revealed significant increases in the anterior cingulate, infralimbic, retrosplenial, auditory, posterior parietal association, perirhinal, and visual areas (FDR < 0.1). In female *Atrx*^NEXCre^ mice, significant increases were observed in the retrosplenial, ectorhinal, posterior parietal association, and visual areas. Conversely, these females exhibited significant decreased volume in the somatomotor, frontal pole, and somatosensory cortical regions (**Figure 3B**).

**Figure 3:**
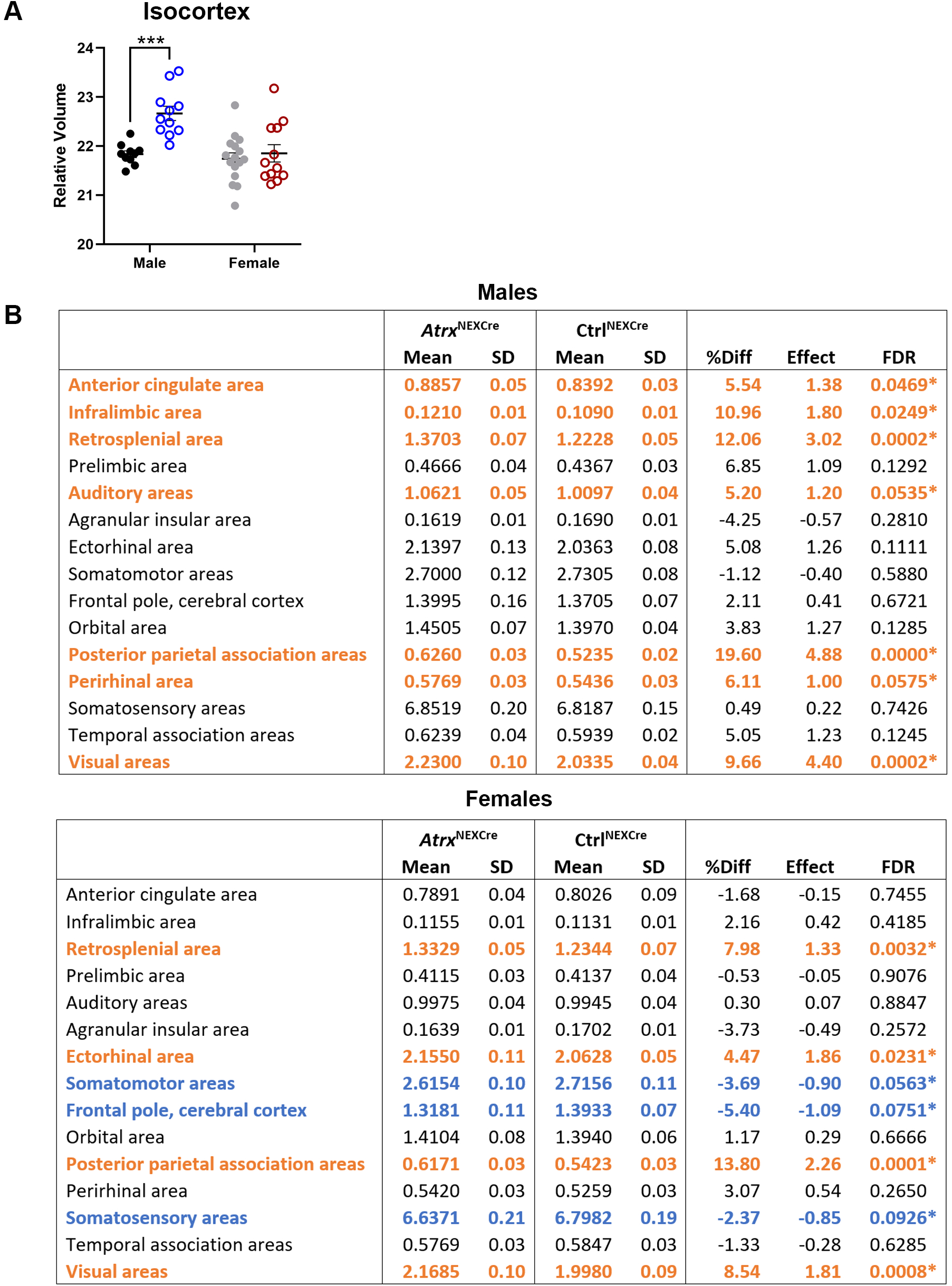
Deletion of ATRX in forebrain excitatory neurons results in relative volume changes in cortical subregions. Tables displaying relative volumes in male and female *Atrx*^NEXCre^ cortical subregions; regions with significantly increased volume are in orange and significantly decreased volume are in orange. SD: standard deviation, FDR: false discovery rate. (Ctrl^male^ n=10, *Atrx*^NEXCre^ ^male^ n=11, Ctrl^female^ n=16, *Atrx*^NEXCre^ ^female^ n=12).

The most pronounced increases in relative volume, observed in both sexes, were localized to the retrosplenial cortex (male FDR = 0.0002, female FDR = 0.0032), posterior parietal association area (male FDR = 0.0002, female FDR = 0.0008), and visual cortex (male FDR < 0.0001, female FDR = 0.0001) (**Figure 4A**). The analysis of subcategories within these areas revealed significant volume increases in cingulate areas 29a and 29b, lateral and medial parietal association areas, the secondary visual cortex, primary visual area, posteromedial visual area, and the anteromedial visual area (**Figure 4B**).

**Figure 4:**
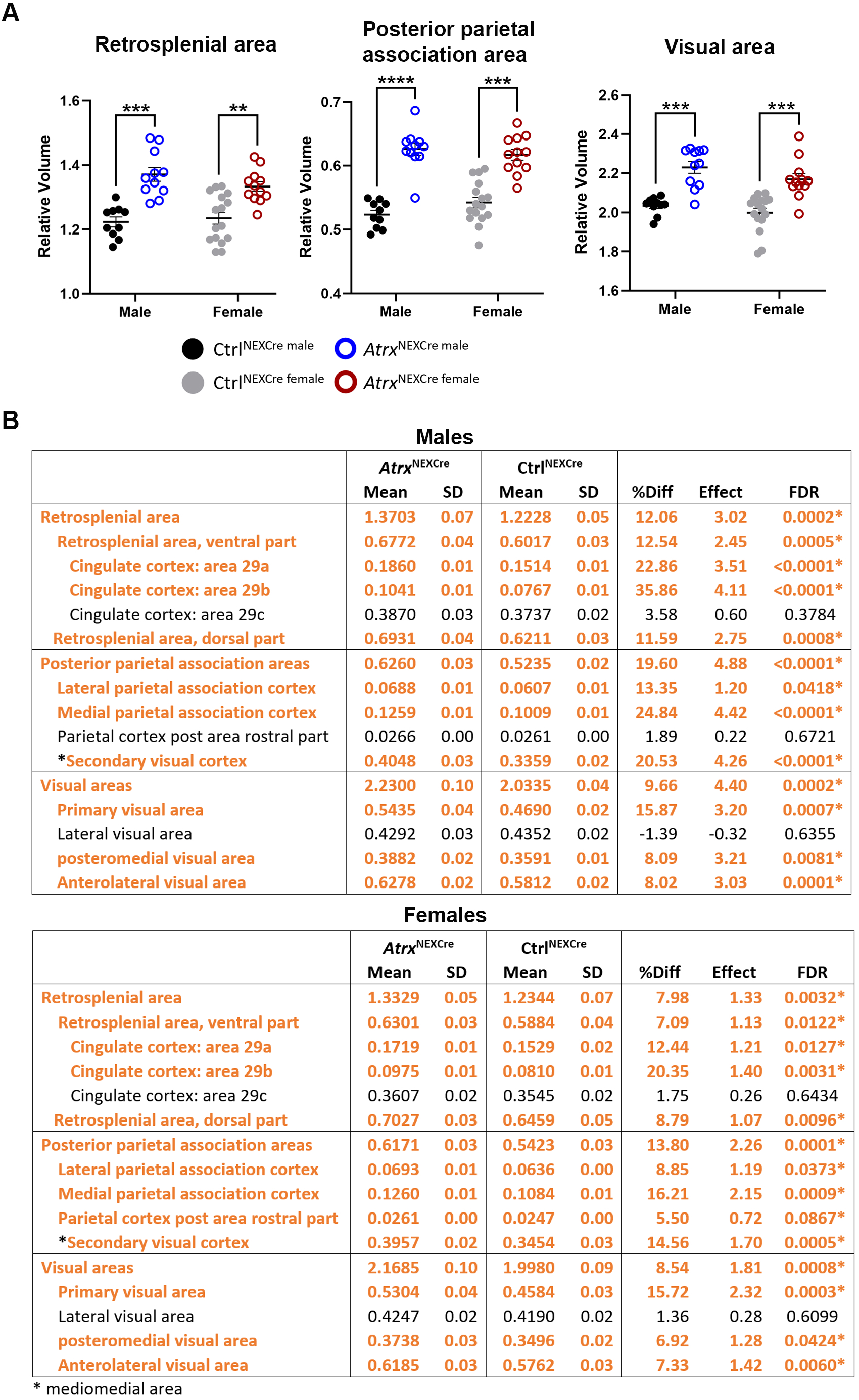
Most enlarged cortical regions in *Atrx*^NEXCre^ mice. **A)** Relative volumes of the most significantly increased isocortex subregions in male and female *Atrx*^NEXCre^ mice. **B)** Tables with breakdown of the retrosplenial, posterior parietal association, and visual area subregions in male and female mice. Orange= regions with significantly increased volume. SD: standard deviation, FDR: false discovery rate.(**= FDR < 0.05, ***= FDR < 0.001, ****= FDR < 0.0001)(Ctrl^male^ n=10, *Atrx*^NEXCre^ ^male^ n=11, Ctrl^female^ n=16, *Atrx*^NEXCre^ ^female^ n=12).

Collectively, these findings demonstrate that ATRX deficiency leads to distinct and region-specific ultrastructural changes in the hippocampus and cortex, with both shared and sex-specific patterns of alteration.

### Structural alterations in brain regions retaining ATRX expression

In addition to regions with ATRX ablation, we observed significant structural changes in multiple brain areas where ATRX expression is retained. Specifically, *Atrx*^NEXCre^ mice had significantly enlarged striatum (male FDR = 0.0001, female FDR = 0.0833), thalamus (male FDR = 0.0002, female FDR = 0.0005), and midbrain (male FDR = 0.0418, female FDR = 0.0025) compared to control mice (**Figure 5A–C**). The hypothalamus was significantly larger only in female *Atrx*^NEXCre^ mice (FDR = 0.0628), while no significant volume changes were detected in the hindbrain in mice of either sex (**Figure 5D–E**).

**Figure 5:**
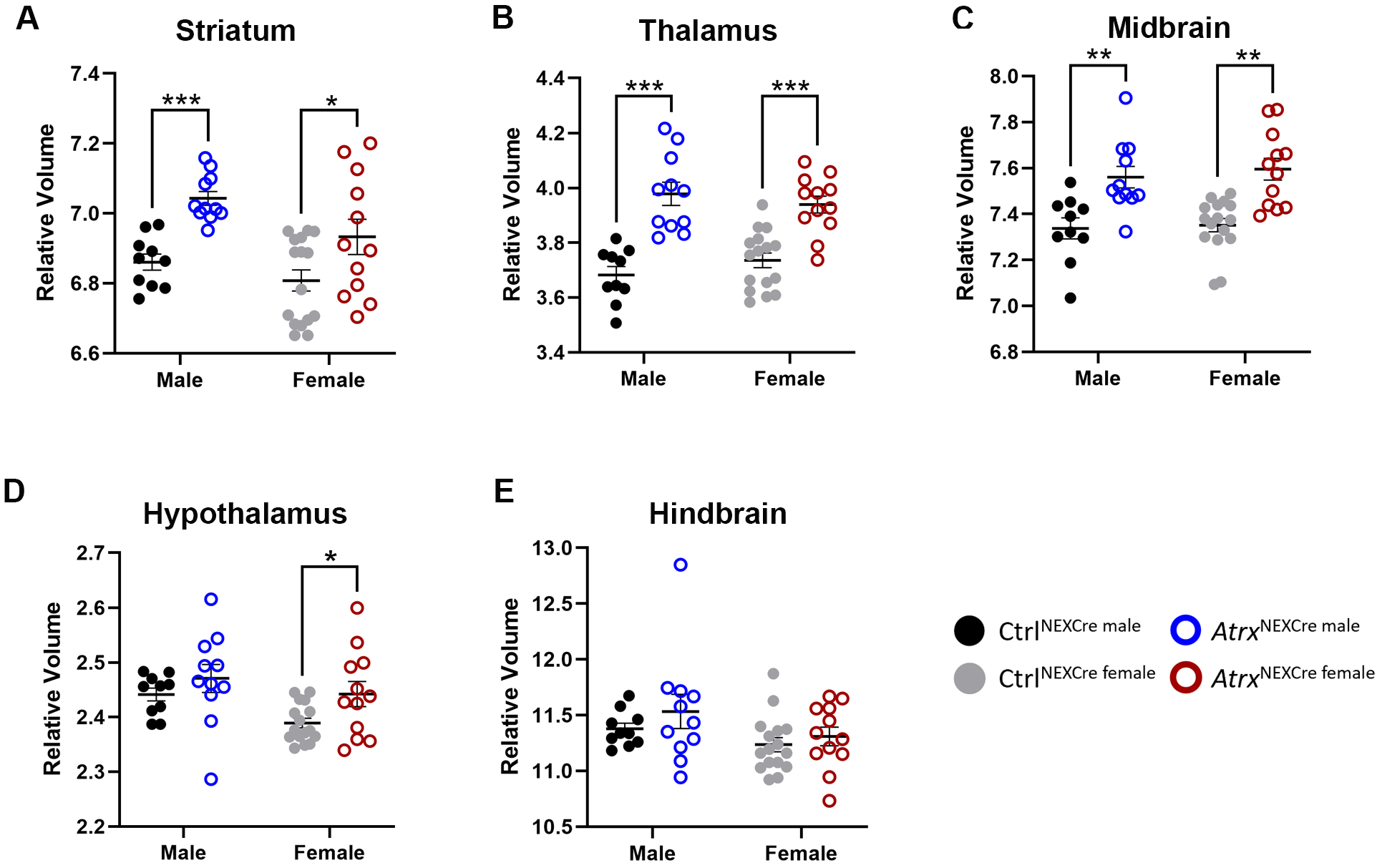
Relative volumetric changes in brain regions that retain ATRX expression in *Atrx*^NEXCre^ mice. Graphs displaying relative brain volumes of **A)** Striatum, **B)** Thalamus, **C)** Midbrain, **D)** Hypothalamus, and **E)** Hindbrain regions in male and female mice. (Ctrl^male^ n=10, *Atrx*^NEXCre^ ^male^ n=11, Ctrl^female^ n=16, *Atrx*^NEXCre^ ^female^ n=12) (*= FDR<0.10, **=FDR<0.05, ***= FDR<0.001).

Multiple cerebellar subregions also displayed structural alterations. These included the cerebellar cortex, arbor vitae (white matter), and the deep cerebellar nuclei, which are critical for integrating and relaying cerebellar input and output signals (Leon & Das, 2023). Notably, the fastigial nucleus showed an increased relative volume in female *Atrx*^NEXCre^ mice (**Figure 6A**).

**Figure 6:**
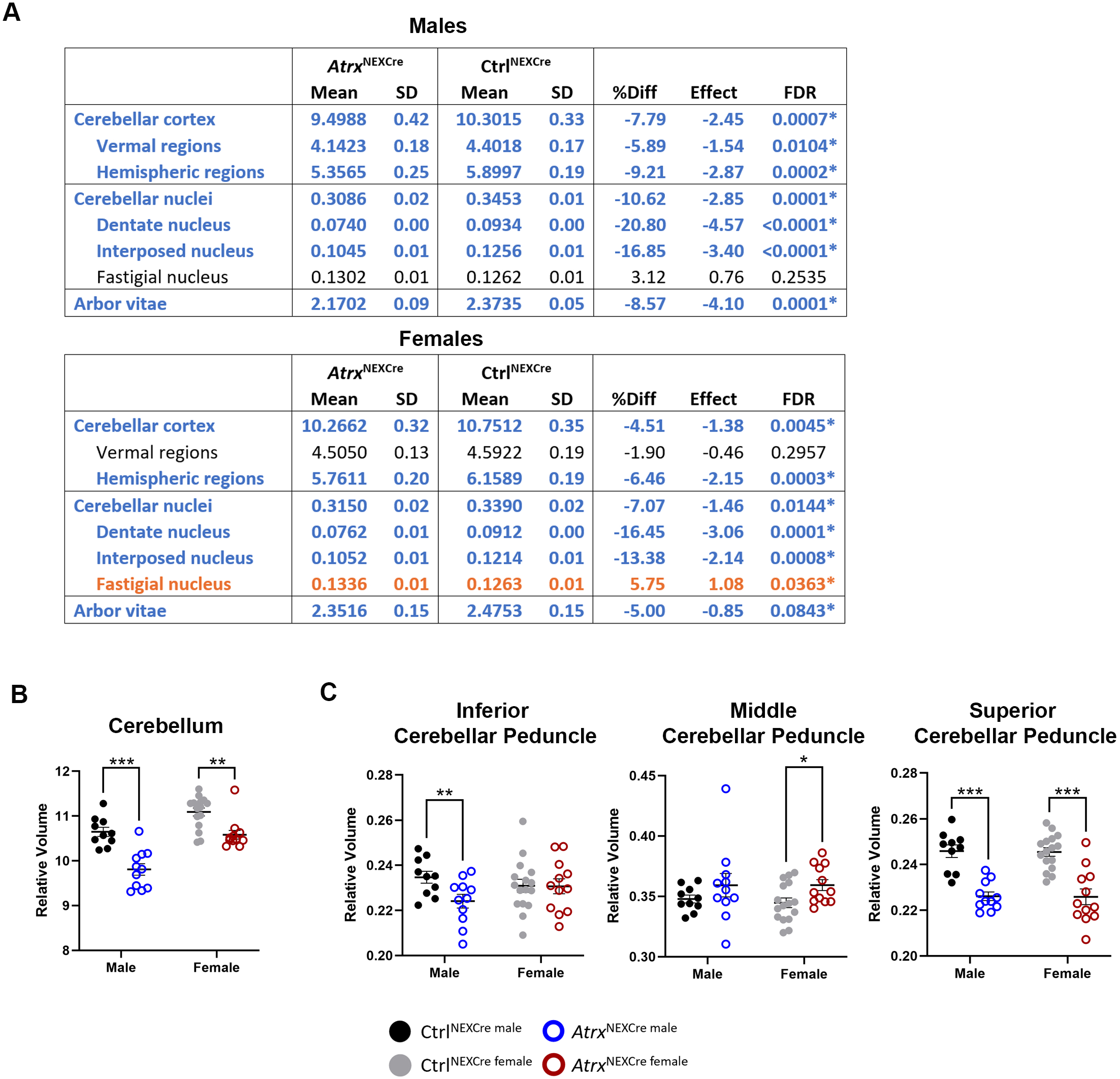
Decreased relative volume of the cerebellum and cerebellar fibre tracts. **A)** Table displaying decrease in the relative brain volume of cerebellum subregions; orange= regions with significantly relative increase in volume and blue= significant relative decrease in volume. SD: standard deviation, FDR: false discovery rate. Graph displaying relative volume changes in **B)** total cerebellum and **C)** cerebellar peduncles in *Atrx*^NEXCre^ mice. Ctrl^male^ n=10, *Atrx*^NEXCre^ ^male^ n=11, Ctrl^female^ n=16, *Atrx*^NEXCre^ ^female^ n=12) (*=FDR <0.10, **=FDR <0.05, ***= FDR < 0.001).

Despite these localized increases, total cerebellar volume was significantly reduced in both males (FDR = 0.0005) and females (FDR = 0.0037) (**Figure 6B**). This reduction was accompanied by alterations in cerebellar fiber tracts: the inferior cerebellar peduncle was significantly decreased in males (FDR = 0.0469), whereas the middle cerebellar peduncle was increased in females (FDR = 0.0618). Importantly, the superior cerebellar peduncle—a major pathway connecting the cerebellum to the midbrain and relaying signals through the dentate nucleus to the ventrolateral thalamic nuclei and red nucleus (Leon & Das, 2023) was significantly decreased in both male (FDR = 0.0001) and female mutant mice (FDR = 0.0003) (**Figure 6C**). ATRX immunofluorescence analysis of *Atrx*^NEXCre^;SUN1-GFP Cre reporter brain sections confirmed that all of the aforementioned regions do not express Cre recombinase and maintain ATRX expression (**Supplemental Figure 1**). These unexpected structural changes in regions retaining ATRX suggest that ATRX deficiency in the developing forebrain can exert indirect, non-cell-autonomous effects on the morphology of interconnected brain regions.

### Bidirectional effects of ATRX deficiency on fibre tracts

In addition to abnormalities in cerebellar-associated fiber tracts, we observed significant alterations in major forebrain fiber tracts in *Atrx*^NEXCre^ mice. The corpus callosum, a critical structure for interhemispheric communication, showed a significant reduction in relative volume in both male (FDR = 0.0006) and female (FDR = 0.0033) *Atrx*^NEXCre^ mice compared to controls. Similarly, the fornix system, which is essential for hippocampal connectivity, was significantly reduced in both males (FDR = 0.0046) and females (FDR = 0.0057) (**Figure 7A-B**). However, not all fiber tracts were reduced in volume. The temporal limb of the anterior commissure exhibited a significant increase in relative volume in males (FDR = 0.0187), while the olfactory limb of the anterior commissure was significantly increased in both sexes (male FDR = 0.0001, female FDR = 0.0001). Additionally, the posterior commissure showed an increased volume in females (FDR = 0.0867) (**Figure 7C-E**). Histological analysis further supported these findings. Examination of brain sections confirmed a significant reduction in the thickness of the corpus callosum at 3 months in both sexes (**Figure 7F**). Collectively, these results demonstrate that ATRX deficiency leads to widespread alterations in structural connectivity, affecting both commissural and projection fiber tracts. The observed changes are largely consistent across sexes, suggesting a fundamental role for ATRX in the development and maintenance of neural connectivity.

**Figure 7:**
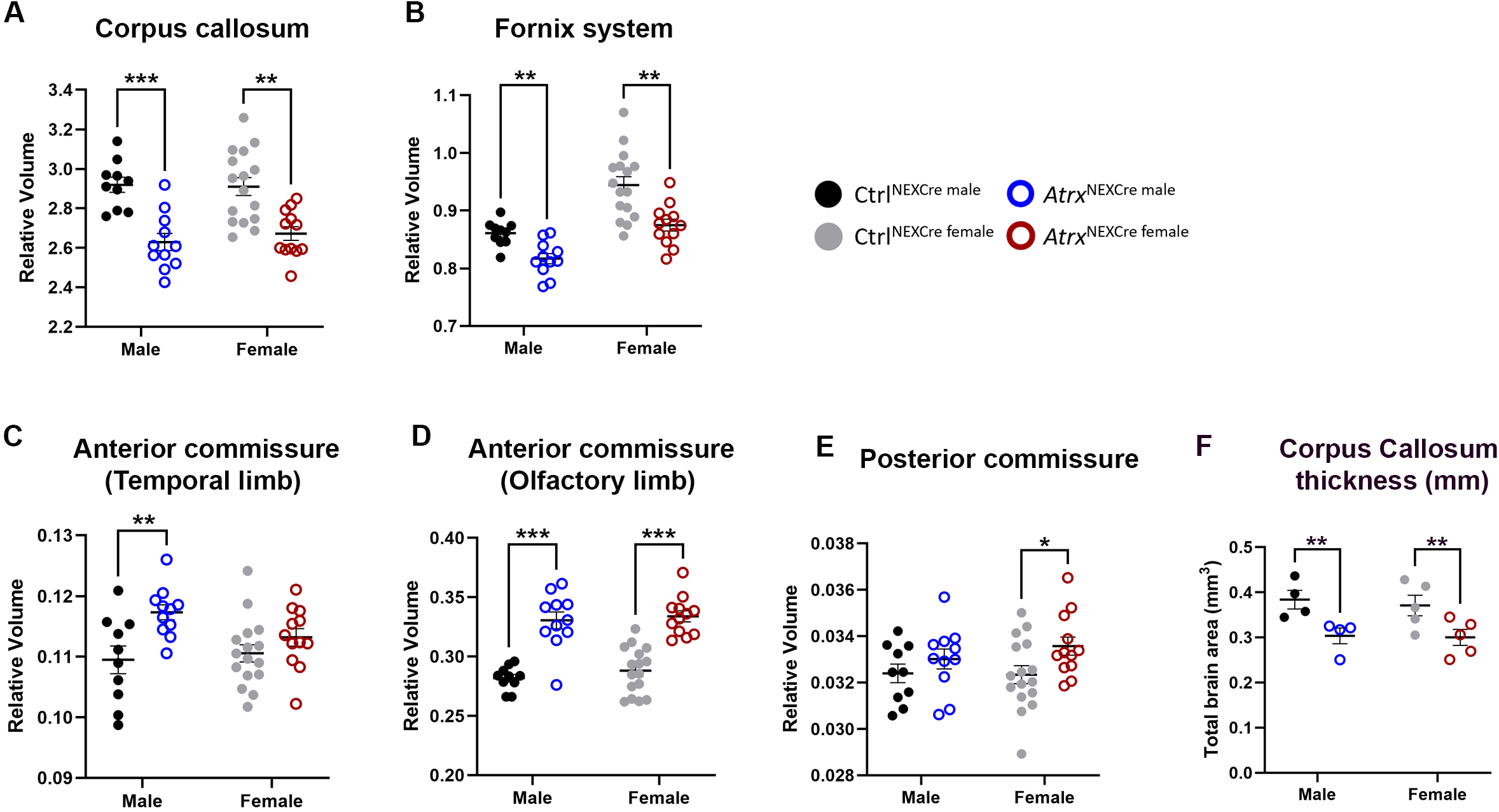
Bimodel effects of ATRX loss on fibre tracts. Graphs displaying decrease in the relative brain volume of **A)** corpus callosum and **B)** fornix system. Increased relative volume of **C)** anterior commissure temporal limb, **D)** olfactory limb, and **E)** posterior commissure. Ctrl^male^ n=10, *Atrx*^NEXCre^ ^male^ n=11, Ctrl^female^ n=16, *Atrx*^NEXCre^ ^female^ n=12. *= FDR <0.10, **=FDR <0.05, *** = FDR< 0.001. F) Measurement of corpus callosum thickness (mm) in histological sections at 3-months (Male n=4, Female n=5) (**=P<0.05).

## DISCUSSION

Our study demonstrates that the loss of ATRX in forebrain excitatory neurons leads to widespread and region-specific structural alterations in the mouse brain. Both male and female *Atrx*^NEXCre^ mice exhibit increased relative volumes in caudal cortical regions, as well as in the thalamus, striatum, and midbrain. At the same time, these mice show reduced volumes in the hippocampus, cerebellum, and associated fiber tracts (**Figure 8**). These findings suggest that ATRX deficiency disrupts not only specific brain regions but also global neural connectivity, which may underlie the cognitive and autism-related behaviors observed in this model.

**Figure 8:**
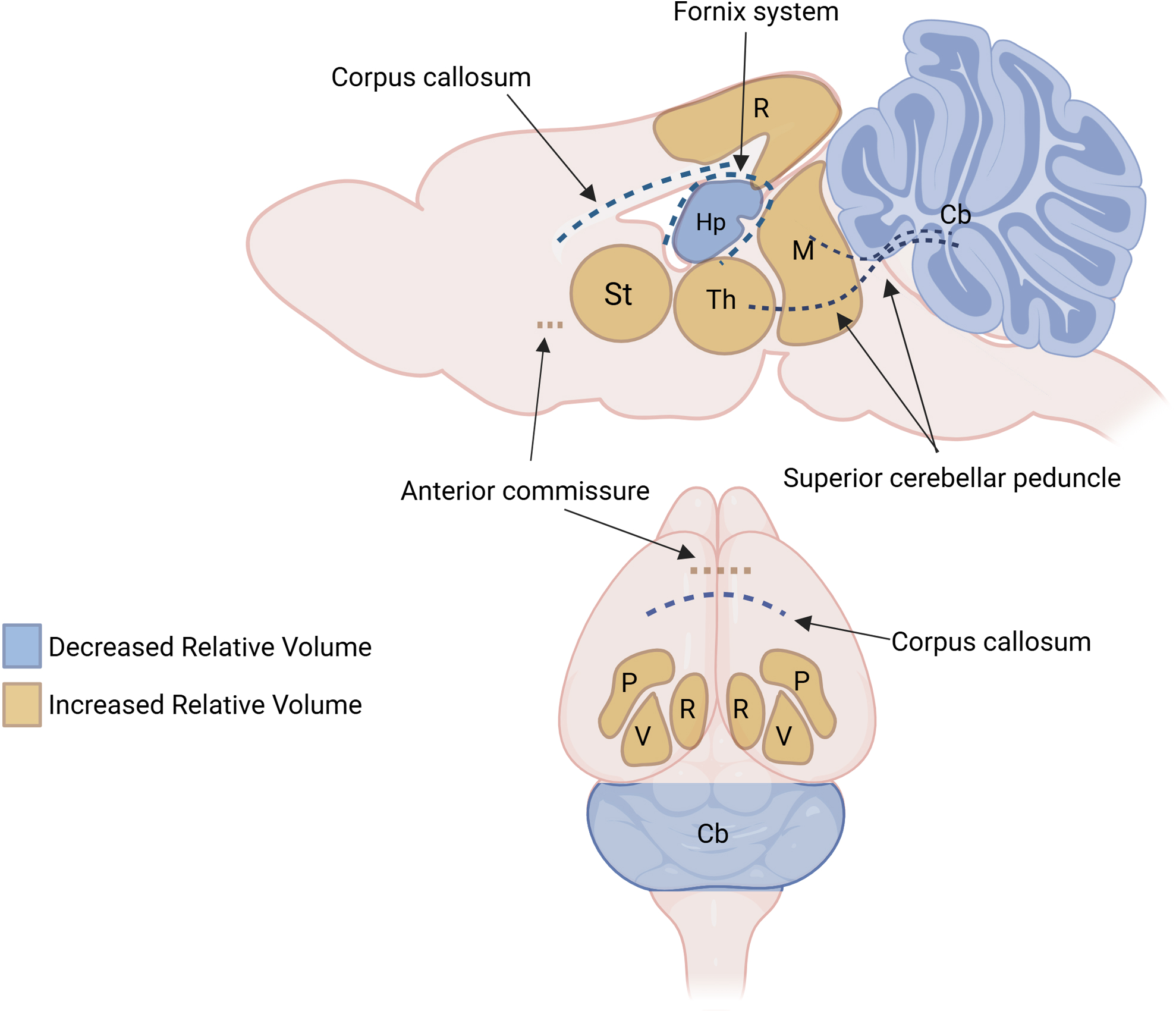
Summary of relative brain volume changes shared by male and female *Atrx*^NEXCre^ mice. Cb= cerebellum, Hp= hippocampus, M= midbrain, P= posterior parietal area, R= retrosplenial area, St= striatum, Th= thalamus, V= visual area.

MRI analysis revealed that male *Atrx*^NEXCre^ mice develop microcephaly, mirroring the microcephaly observed in ATR-X syndrome patients (Gibbons & Higgs, 2000). Although mutant mice of both sexes displayed reduced body size, only males exhibited a significant reduction in total brain volume. This pattern suggests a sex-dependent vulnerability to ATRX loss. The underlying cause of these growth differences remains unclear, but similar findings have been reported in other mouse models with ATRX deletion in the central nervous system (Tamming et al., 2017, 2020). The reduced hippocampal volume and fornix observed in *Atrx*^NEXCre^ mice are consistent with memory and cognitive deficits, which are also reported in both this mouse model and ATR-X syndrome (Gibbons & Higgs, 2000; Quesnel et al., 2023). These findings are similar to those seen in other mouse models with cognitive deficits, including models with mutations in *Nlgn3*, an ASD-associated gene encoding a synaptic adhesion protein, and *MeCP2*, a gene encoding a chromatin remodeling protein associated with Rett syndrome (Akaba et al., 2022; Anand & Dhikav, 2012; Ellegood et al., 2011, 2025). However, since models with postnatal *Atrx* deletion show hippocampal deficits without autism features (Quesnel et al., 2023; Martin-Kenny & Bérubé, 2020), these changes may be more closely linked to intellectual disability than to autism itself.

Our analysis also reveals significantly increased volumes in the retrosplenial, posterior parietal, and visual cortical areas of the caudal cortex in *Atrx*^NEXCre^ mice. In contrast, reductions in these same regions have been reported in ASD mouse models with MeCP2 or *Chd7* mutations, suggesting that volumetric alterations in these cortical areas are linked to ASD-associated behaviors (Akaba et al., 2022; Donovan et al., 2023). The retrosplenial cortex has been implicated in social memory deficits and repetitive behaviors (Garrido et al., 2022; Ha et al., 2015; Thakkar et al., 2008), while the posterior parietal cortex is involved in spatial awareness and working memory (Hahn et al., 2018; Sack, 2009). Consistent with these roles, *Atrx*^NEXCre^ mice display cognitive and social memory deficits, as well as repetitive behaviors (Quesnel et al., 2023). These results suggest that structural changes in these caudal cortical regions may contribute directly to the behavioral phenotypes observed with ATRX deficiency.

Alterations in the visual cortex are frequently associated with sensory processing deficits in ASD, including difficulties interpreting visual cues such as facial expressions and body language. Although visual processing has not yet been specifically assessed in *Atrx*^NEXCre^ mice, MeCP2—a chromatin-associated protein that interacts with ATRX—modulates visual cortical neuron activity, and its loss reduces visual activity in mice (Chung & Son, 2020; Zhang et al., 2017). We also observed sex-specific differences in cortical structure. For example, males showed increased auditory cortex volume and related sensory processing deficits, such as increased auditory startle response and impaired auditory sensory gating, which were not observed in female mice (Quesnel et al., 2023). These results indicate that structural changes in sensory regions may underlie sex differences in autism-related behaviors, consistent with clinical observations of ASD.

Interestingly, we identified structural alterations in brain regions that retain ATRX expression, indicating that ATRX deficiency in forebrain excitatory neurons can have non-cell-autonomous effects, potentially through disrupted connectivity or altered developmental signaling. Alterations in major fiber tracts, including the corpus callosum and fornix, further highlight disrupted connectivity in *Atrx*^NEXCre^ mice.

Our findings support the idea that cerebellar alterations could underlie ASD behaviors. The cerebellum has functional connectivity to the cortex, hippocampus, and amygdala, and is important for cognitive and emotional function. Defects in cerebellar structures are reported in neurodevelopmental disorders, including ASD, ADHD, and intellectual disability (Mapelli et al., 2022; Sathyanesan et al., 2019). In addition, pediatric patients with cerebellar lesions often exhibit intellectual disability and ASD symptoms, including language deficits, repetitive behaviors, and decreased sociability (Hampson & Blatt, 2015). The cerebellum has extensive “closed-loop” connections, forming specific circuits with regions of the cortex and may act as an integrative processor for input and output responses (Courchesne et al., 2007; D’Mello & Stoodley, 2015). The most significant reduction in relative brain volume in *Atrx*^NEXCre^ mice was observed in the dentate nucleus, a deep cerebellar nucleus involved in connecting the cerebellum with the forebrain, a finding also seen in many other ASD mouse models (Ellegood et al., 2015). The superior cerebellar peduncle, which projects out of the cerebellum through the dentate nucleus to the midbrain and thalamus, also exhibited a significant reduction in relative volume in *Atrx*^NEXCre^ mice (Hampson & Blatt, 2015; Leon & Das, 2023). A previous study in mice demonstrated that disruption of cortico-thalamic-cerebellar connectivity results in ASD features (Kelly et al., 2020). Taken together, the alterations in relative volumes identified in *Atrx*^NEXCre^ mice are consistent with aberrant cortical-thalamic-cerebellar connectivity.

Early fiber tract alterations in individuals with ASD have been proposed to contribute to the atypical development of brain connectivity (Ha et al., 2015). In this study, MRI data revealed a reduced volume of the corpus callosum in *Atrx*^NEXCre^ mice, a hallmark in the brains of individuals with ASD and schizophrenia (Domínguez-Iturza et al., 2019; Saxena et al., 2012) and ASD/intellectual disability mouse models (Akaba et al., 2022; Donovan et al., 2023; Ellegood et al., 2011). Theories of structural covariance suggest that brain regions are both structurally and functionally connected, potentially due to coordinated growth driven by the expression of shared gene sets during development. The cortical-thalamic relay of information through structural and functional connection is an example of structural covariance (Yee et al., 2024). Disruption of developmental gene expression patterns due to loss of ATRX in forebrain excitatory neurons could lead to alterations in both structural connectivity and the gene expression profiles required for the growth and maturation of subcortical regions such as the thalamus and striatum. Further investigation is needed to characterize the specific gene expression changes in *Atrx*^NEXCre^ mice during development and how these alterations might contribute to impaired subcortical development.

This study is limited by its focus on volumetric MRI, which cannot directly assess functional or structural connectivity. Future work using functional MRI and diffusion tensor imaging will be important to clarify the impact of ATRX loss on brain networks. Additionally, examining these changes across development could identify critical periods for intervention. Further investigation into the mechanisms driving regional volume increases, particularly in sensory cortices, may provide insight into sex-specific sensory processing deficits in ASD and inform future therapeutic strategies.

In summary, our findings highlight the essential role of ATRX in shaping brain structure and connectivity. The observed alterations in cortical, hippocampal, cerebellar, and subcortical regions in *Atrx*^NEXCre^ mice mirror key features of ASD and intellectual disability. These results support the use of this model to study the neural basis of autism and related neurodevelopmental disorders.

## DECLARATIONS

### Availability of data and materials

All data generated or analyzed during this study are included in this published article.

### Ethics approval and consent to participate

Not applicable

### Author Contributions

K.Q. contributed to conceptualization, design and execution of experiments, data interpretation, and writing of article. J.E and J.P.L. contributed to execution of experiments, data analysis and manuscript editing. N.G.B. contributed the conception, design, interpretation of data, and writing of article.

## FIGURE LEGENDS

**Supplemental Figure 1: Deletion of ATRX and Cre expression is specific to the forebrain.**

**A)** Cre-dependent expression of SUN1-GFP (green) is observed in nuclei of the cortex correlating with loss of ATRX (red) expression. **B)** The striatum, **C)** thalamus, **D)** midbrain and **E)** cerebellum nuclei do not express SUN1-GFP correlating with no loss of ATRX in these brain regions. (Representative images n=3, scale bar= 200um). **F)** Cre-dependent expression of tomato reporter labelled neurons, showing similar expression patterns between Ctrl^NEXCre^ and *Atrx*^NEXCre^ (representative images n=3).

