## Supplementary Figure 1 for "Altered brain structure in an ATRX-deficient mouse model of autism spectrum disorder"

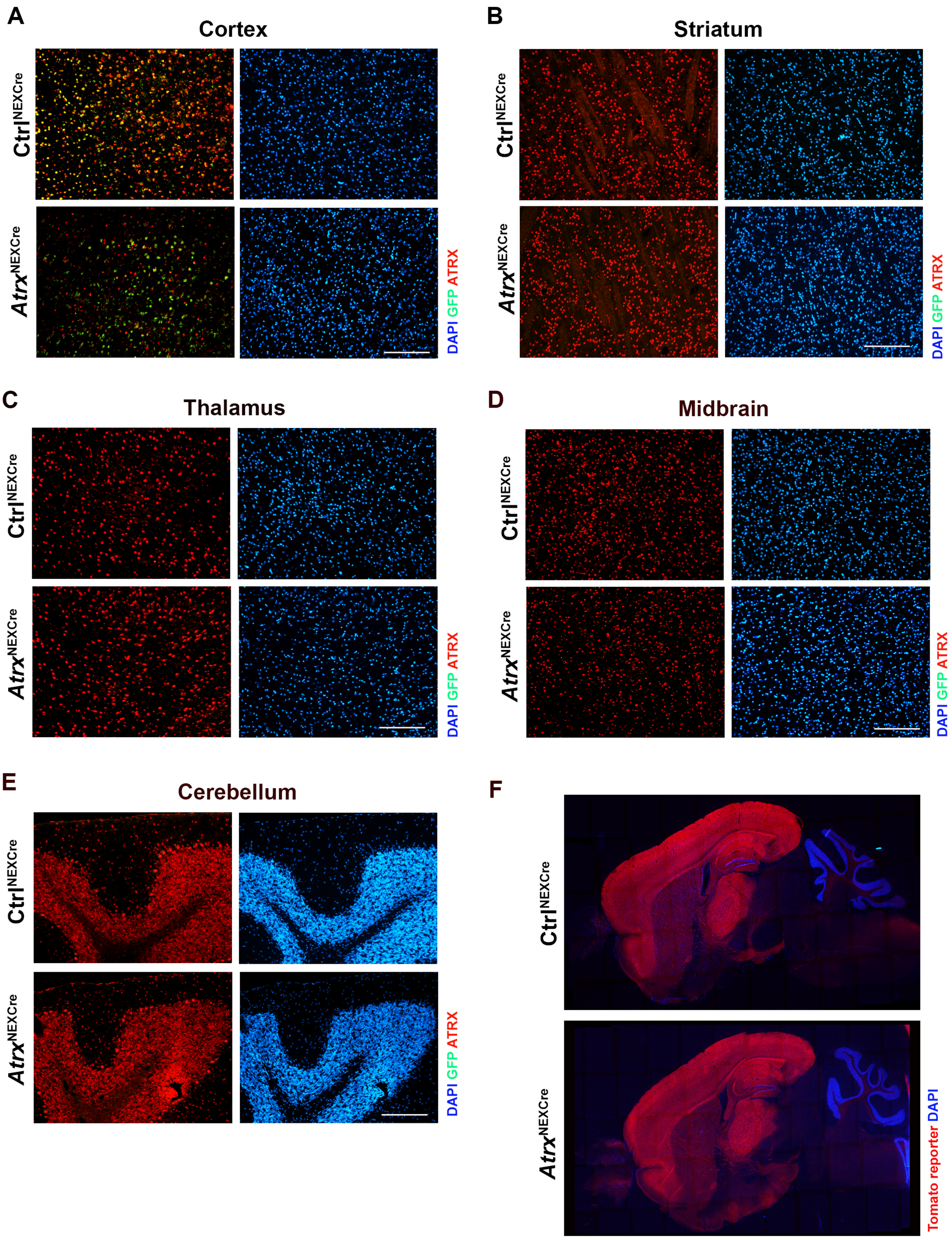


**Supplemental Figure 1: Deletion of ATRX and Cre expression is specific to the forebrain. A)** Cre-dependent expression of SUN1-GFP (green) is observed in nuclei of the cortex correlating with loss of ATRX (red) expression. **B)** The striatum, **C)** thalamus, **D)** midbrain and **E)** cerebellum nuclei do not express SUN1-GFP correlating with no loss of ATRX in these brain regions. (Representative images n=3, scale bar= 200um). **F)** Cre-dependent expression of tomato reporter labelled neurons, showing similar expression patterns between Ctrl$^{NEXCre}$ and *Atrx*$^{NEXCre}$ (representative images n=3).
